# Large-scale transcriptome-wide association study identifies new prostate cancer risk regions

**DOI:** 10.1101/345736

**Authors:** Nicholas Mancuso, Simon Gayther, Alexander Gusev, Wei Zheng, Kathryn L. Penney, Zsofia Kote-Jarai, Rosalind Eeles, Matthew Freedman, Christopher Haiman, Bogdan Pasaniuc

## Abstract

Although genome-wide association studies (GWAS) for prostate cancer (PrCa) have identified more than 100 risk regions, most of the risk genes at these regions remain largely unknown. Here, we integrate the largest PrCa GWAS (N=142,392) with gene expression measured in 45 tissues (N=4,458), including normal and tumor prostate, to perform a multi-tissue transcriptome-wide association study (TWAS) for PrCa. We identify 235 genes at 87 independent 1Mb regions associated with PrCa risk, 9 of which are regions with no genome-wide significant SNP within 2Mb. 24 genes are significant in TWAS only for alternative splicing models in prostate tumor thus supporting the hypothesis of splicing driving risk for continued oncogenesis. Finally, we use a Bayesian probabilistic approach to estimate credible sets of genes containing the causal gene at pre-defined level; this reduced the list of 235 associations to 120 genes in the 90% credible set. Overall, our findings highlight the power of integrating expression with PrCa GWAS to identify novel risk loci and prioritize putative causal genes at known risk loci.

## Introduction

Prostate cancer (PrCa) affects ~1 in 7 men during their lifetime and is one of the most common cancers worldwide, with up to 58% of risk due to genetic factors^1^^;^ ^2^. Genome-wide association studies (GWAS) have identified over 100 genomic regions harboring risk variants for PrCa which explain roughly one third of familial risk^3^^-^^7^. With few exceptions^8^, the causal variants and target susceptibility genes at most GWAS risk loci have yet to be identified. Multiple studies have shown that PrCa- and other disease-associated variants are enriched near variants that correlate with gene expression levels^9^^-^^13^. In fact, recent approaches have integrated expression quantitative trait loci (eQTLs) with GWAS to implicate several plausible genes for PrCa risk (e.g., *IRX4, MSMB, NCOA4, NUDT11* and *SLC22A3)*^5^^;^ ^14^^-^^21^. While overlapping eQTLs and GWAS is powerful, the high prevalence of eQTLs^22^ coupled with linkage disequilibrium (LD) renders it difficult to distinguish the true susceptibility gene from spurious co-localization at the same locus^23^. Therefore, disentangling LD is critical for prioritization and causal gene identification at risk loci.

Gene expression imputation followed by a transcriptome-wide association study^24^^-^^26^ (TWAS) has been recently proposed as a powerful approach to prioritize candidate risk genes underlying complex traits. By taking LD into account across SNPs, the resulting association statistics reflect the underlying effect of steady-state gene or alternative splicing expression levels on disease risk^25^^;^ ^27^, which can be used to identify new regions or to rank genes for functional validation at known risk regions^24^^-^^28^. Here we perform a multi-tissue transcriptome-wide association study^24^^-^^26^ to identify new risk regions and to prioritize genes at known risk regions for PrCa. Specifically, we integrate gene expression data from 48 panels measured in 45 tissues across 4,448 individuals with GWAS of prostate cancer from the OncoArray in 142,392 men^29^. Notably, we include alternatively spliced and total gene expression data measured in tumor prostate to identify genes contributing to prostate cancer risk or to continued oncogenesis. We identify 235 gene-trait associations for PrCa with 24 (11) genes identified uniquely using models of alternative spliced (total) expression in tumor. Significant genes were found in 87 independent 1Mb regions, of which 9 regions are located more than 2Mb away from any OncoArray GWAS significant variants, thus identifying new candidate risk regions. Second, we use TWAS to investigate genes previously reported as susceptibility genes for prostate cancer identified by eQTL-based analyses. We find a significant overlap with 57 out of 104 previously reported genes assayed in our study also significant in TWAS. Third, we use a novel Bayesian prioritization approach to compute credible sets of genes and prioritize 120 genes that explain at least 90% of the posterior density for association signal at TWAS risk regions. One notable example, *IRX4,* had 97% posterior probability to explain the association signal at its region with the remaining 3% explained by 9 neighboring genes. Overall, our findings highlight the power of integrating gene expression data with GWAS and provide testable hypotheses for future functional validation of prostate cancer risk.

## Results

### Overview of methods

To identify genes associated with PrCa risk, we performed a TWAS using 48 gene expression panels measured in 45 tissues^22^^;^ ^30^^-^^36^ integrated with summary data from the OncoArray PrCa GWAS of 142,392 individuals of European ancestry (81,318/61,074 cases/controls; see Methods)^29^. We performed the summary-based TWAS approach as described in ref^25^ using the FUSION software (see Methods). Briefly, this approach uses reference linkage-disequilibrium (LD) and reference gene expression panels with GWAS summary statistics to estimate the association between the cis-genetic component of gene expression, or alternative splicing events, and PrCa risk^25^. First, for each panel, FUSION estimated the heritability of steady-state gene and alternative splicing expression levels explained by SNPs local to each gene (i.e. 1Mb flanking window) using the mixed-linear model (see Methods). Genes with nominally significant (*P* < 0.05) estimates of SNP-heritability (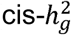), are then put forward for training predictive models. Genes with non-significant estimates of heritability are pruned, as they are unlikely to be accurately predicted. Next, FUSION fits predictive linear models (e.g., Elastic Net, LASSO, GBLUP^37^, BSLMM^38^) for every gene using local SNPs. The model with the best cross-validation prediction accuracy (out-of-sample *R*^2^) was used for prediction into the GWAS cohort. This was repeated for all expression datasets, resulting in 117,459 tissue-specific models spanning 16,052 unique genes using total expression and 5,140 using alternatively spliced introns for a combined 17,023 unique genes. The average number of models per expression panel was 2397.6 (see Table S1). Gene expression measured in normal prostate tissue from GTEx^22^ resulted in only 854 gene models, which can be explained due to smaller sample size (*N* = 87) compared with the average (*N* = 234; see Table S1). Indeed, the number of gene models per panel was highly correlated with sample size, which implies that statistical power to detect genes with cis-regulatory control is limited by sample size (see Figure S1). Focusing only on models capturing total gene expression, genes on average had heritable levels of expression in 6.4 different panels (median 3) with 11,364 / 16,052 genes having heritable expression in at least 2 panels (see Figure 1). Predictive power of linear gene expression models is upper-bounded by heritability; thus, we use a normalized *R*^2^ to measure in-sample prediction accuracy (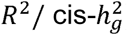). We found the average 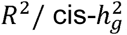 across all tissue-specific models was 61%, which indicates that most of the signal in cis-regulated total expression and alternative splicing levels is captured by the fitted models (see Figure 1). To assess the predictive stability for models of normal prostate gene expression, we compared measured and predicted gene expression for TCGA^36^^;^ ^39^ samples using models fitted in GTEx^22^ normal prostate. We found a highly significant replication (*R*^2^ = 0.07; *P =* 1.5 × 10^−29^), explaining 39% of in-sample cross-validation *R*^2^ (see Figure S2), which is consistent with previous out-of-sample estimates^24^^;^ ^25^. We performed a cross-tissue analysis within TCGA and found tumor prostate gene expression models replicated in normal prostate (total expression *R*^2^ = 0.06; splicing *R*^2^ = 0.05; see Table S2). Given the large number of genes having evidence of genetic control across multiple tissues, we next aimed to measure the similarity of different tissue models (see Methods). Across all reference panels for each gene we observed an average *R*^2^ = 0.64 (see Figure S3). Similarly, when averaging across genes, reference panels displayed an average cross-tissue *R*^2^ = 0.52 (see Figure S4). Together, these results suggest that trained models predict similar levels of cis-regulated expression on average, despite reference panels measuring expression in different tissues, from varying QC, and capture technologies. Next, we performed simulations to measure the statistical power of TWAS under a variety of trait architectures (see Supplementary Note). Consistent with previous work, we found TWAS to be well-powered at various effect-sizes and heritability levels for gene expression. Importantly, we found no inflation under the null when cis-regulated gene expression has no effect on downstream trait (see Figure S5).

**Figure 1.**
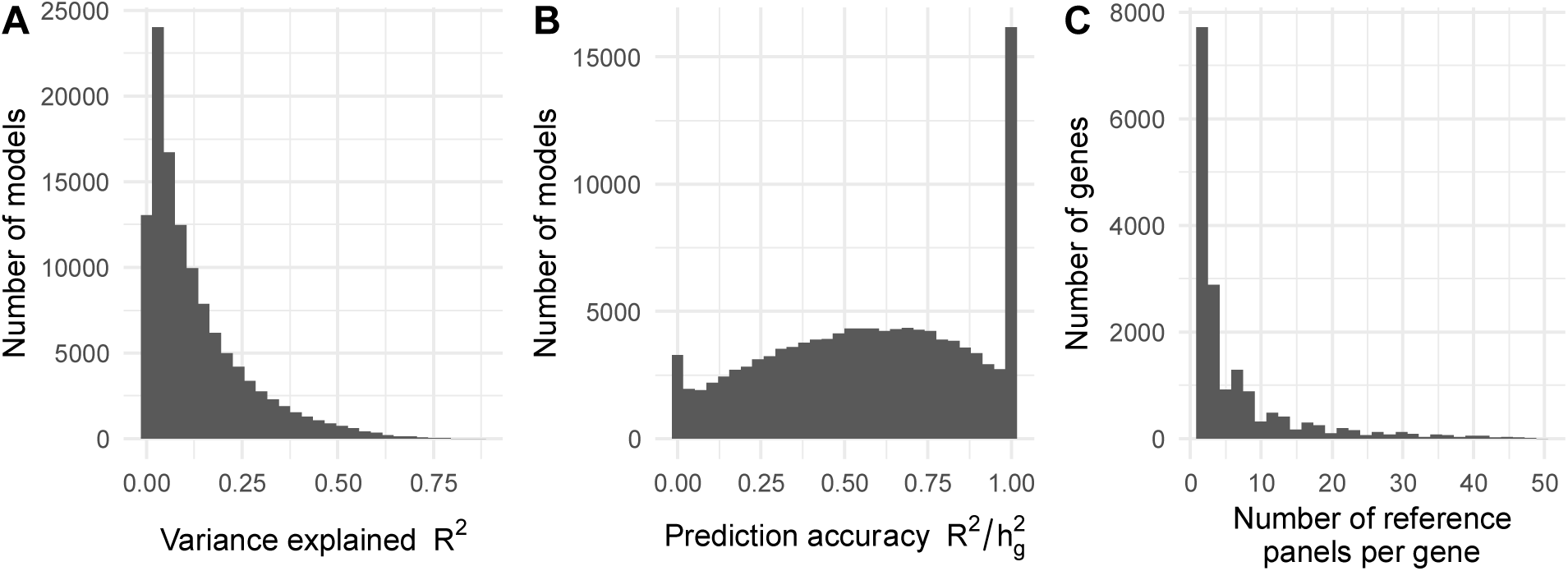
Tissue-specific predictive models for gene expression. A) Cross-validation prediction accuracy of cis-regulated expression and splicing events (*R*^2^) for all 117,459 tissue-specific models. B) Normalized prediction accuracy (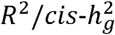) for all 117,459 tissue-specific models. C) Histogram of the number of reference panels per gene. The majority of genes were heritable in a small number of tissues, but many genes exhibited heritable levels across many tissues.

### Multi-tissue TWAS identifies 235 genes associated with PrCa status

In total, we tested 117,459 tissue-specific gene models of expression for association with PrCa status and observed 932 reaching transcriptome-wide significance (*P_TWAS_* < 4.26 × 10^−7^), resulting in 235 unique genes, of which 118 were significant in more than one panel (see Table S3; Figure 2). On average, we found 16.8 tissue-specific models associated with PrCa per reference expression panel (see Table S1). In 1Mb regions with at least 1 transcriptome-wide significant gene, we observed 10.7 tissue-specific associated models on average, and 2.7 associated genes on average, indicating that further refinement of association signal at TWAS risk loci is necessary. To quantify the overlap between non-HLA, autosomal risk loci in the OncoArray PrCa GWAS and our TWAS results, we partitioned GWAS summary data into 1Mb regions and observed 131 harboring at least one genome-wide significant SNP. Of these, 126/131 overlapped at least one gene model in our data and 68/131 overlapped at least one transcriptome-wide significant gene (see Figure S6). Associated genes were the closest gene to the top GWAS SNP 20% of the time when using 26,292 RefSeq genes. This result is consistent with previous reports^9^^;^ ^25^^;^ ^26^ and suggests that prioritizing genes based on distance to index SNPs is suboptimal. We found gene model associations were largely consistent, further supporting the predictive stability of models using cis-SNPs (see Figure S7; Supplementary Note). We observed little evidence of prediction accuracy introducing biased results (see Figure S8; Supplementary Note). As a partial control, we compared TWAS results with S-PrediXcan, a related method for predicting gene expression into GWAS summary statistics, using independently trained models and observed a strong correlation (*R* = 0.87; see Figure S9; Supplementary Note), further supporting the validity of the TWAS approach.

**Figure 2.**
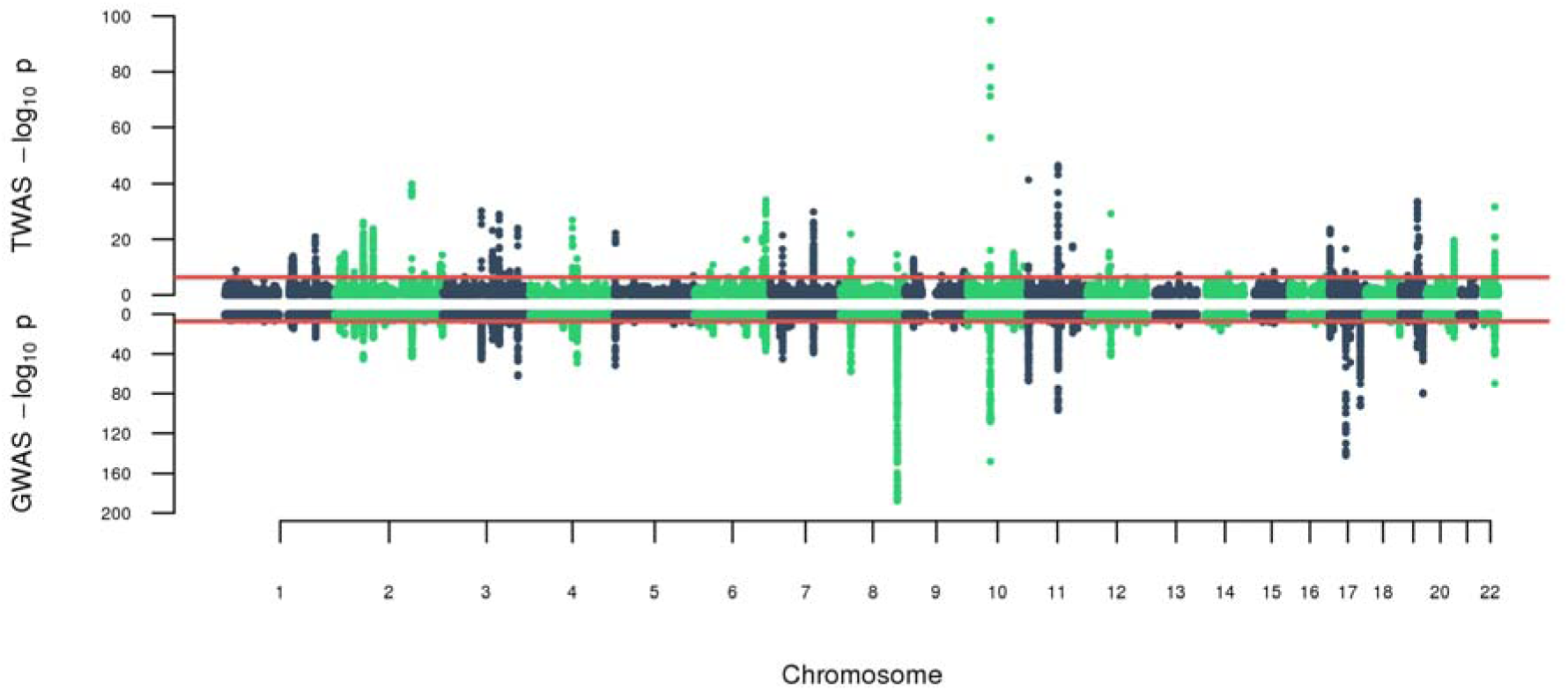
OncoArray PrCa TWAS and GWAS. The top figure is the TWAS Manhattan plot. Each point corresponds to an association test between predicted gene expression with PrCa risk. The red line represents the boundary for transcriptome-wide significance (4.26 × 10^−7^). The bottom figure is the GWAS Manhattan plot where each point is the result of a SNP association test with PrCa risk. The red line corresponds to the traditional genome-wide significant boundary (5 × 10^−8^).

Most of the gene models captured total expression levels in normal tissues, however as a positive control we included models for total expression in tumor prostate tissue (see Methods). Predicted expression using tumor prostate models accounted only for 42/235 significant genes compared with 6/235 in normal prostate which is likely due to the large difference in sample size between the original reference panels (see Table S1). Given this, we found no significant increase in proportion of tumor prostate associated models compared with normal prostate (Fisher’s exact *P* = 0.27). Of the 335 genes with models trained in both reference panels a single shared gene, *MLPH* (OMIM: 606526, a gene whose function is related to melanosome transport^40^), was associated with PrCa risk (see Table S2). 11/42 genes were significant only in tumor prostate models of total expression. 7/11 genes were modeled in other panels but did not reach transcriptome-wide significance while the other 4/11 were not significantly heritable, and thus not testable, in other panels. We also tested models of alternatively spliced introns for association to PrCa risk. We identified predicted expression of alternatively spliced introns in tumor prostate accounted for 69/235 genes, with an average of 2.5 (median 1) alternatively spliced intron associations per gene. We next quantified the amount of overlap between results driven from models of alternative splicing events versus models of total gene expression. 24/69 genes were found only in alternatively spliced introns, and 16/24 genes had models of total gene expression but did not reach transcriptome-wide significance. The remaining 8/24 were tested solely in alternatively spliced introns, due to heritability of total gene expression not reaching significance. Together these results emphasize earlier work demonstrating that sQTLs for a gene commonly capture signal independent of eQTLs^41^.

### TWAS analysis increases power to find PrCa associations

Most of the power in the TWAS approach can be attributed to large GWAS sample size. However, two other factors can increase power over GWAS. First, TWAS carries a reduced testing burden compared with that of GWAS, due to TWAS having many fewer genes compared with SNPs. 10/235 genes were located at 9 novel independent 1Mb regions (i.e. no overlapping GWAS SNP), all of which remained significant under a summary-based permutation test (*P* < 0.05 / 10; see Table 1; Table S2; Methods). We found this result was stable to increasing region sizes (see Table S4) and unlikely be the result of long-range tagging with known GWAS risk (see Table S5; Supplemental Note). We observed increased association signal for SNPs at these regions compared to the genome-wide background after accounting for similar MAF and LD patterns (see Figure S10), which, together with observed TWAS associations, suggests that GWAS sample size is still a limiting factor in identifying PrCa risk SNPs. As a partially independent check, we performed a multi-tissue TWAS using summary data from an earlier PrCa GWAS (*N* = 49,346)^7^ and found 2 novel regions. We found both regions to overlap a genome-wide significant SNP within 1Mb in this data further supporting the robustness of TWAS (see Table S6). Second, we expect to observe increased association signal when expression of a risk gene is regulated by multiple local SNPs^25^. We observed 90/932 instances across 31 genes where TWAS association statistics were stronger than the respective top overlapping GWAS SNP statistics (one-sided Fisher’s exact *P* < 2.2 × 10^−16^; 7% higher *χ*^2^ statistics on average). For example, *GRHL3* (OMIM:608317; a gene associated with suppression of squamous cell carcinoma tumors^42^) exhibited stronger signal in TWAS using expression in prostate tumor (*P_TWAS_* = 9.38 × 10^−10^) compared with the lead SNP signal (*P_GWAS_ =* 1.49 × 10^−5^). Similarly, *POLI* (OMIM:605252, a DNA repair gene associated with mutagenesis of cancer cells^43^^;^ ^44^) resulted in larger TWAS associations (*P_TWAS_* = 2.01 × 10^−7^) compared with the best proximal SNP (*P_GWAS_* = 5.44 × 10^−7^).

**Table 1.**
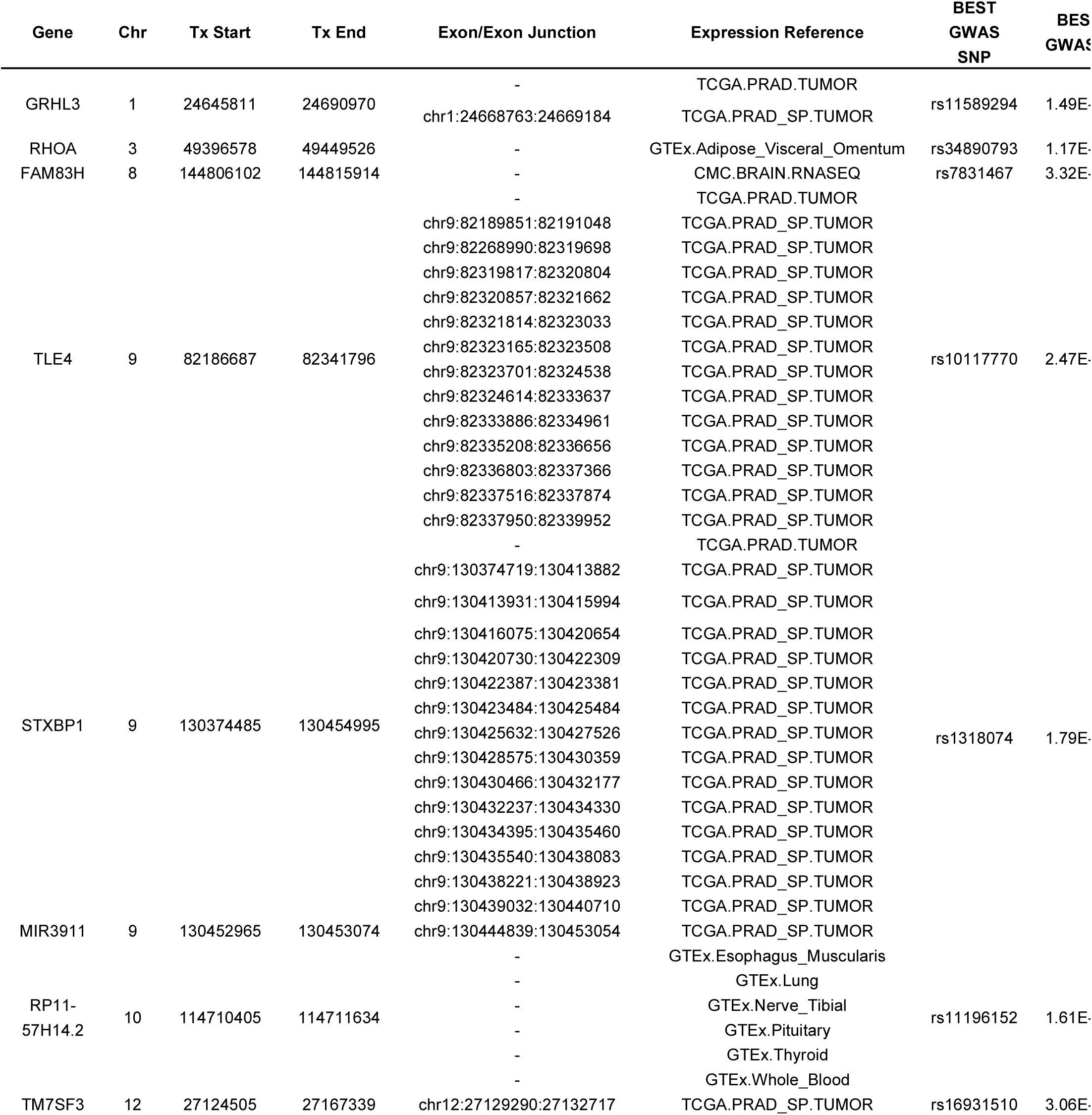
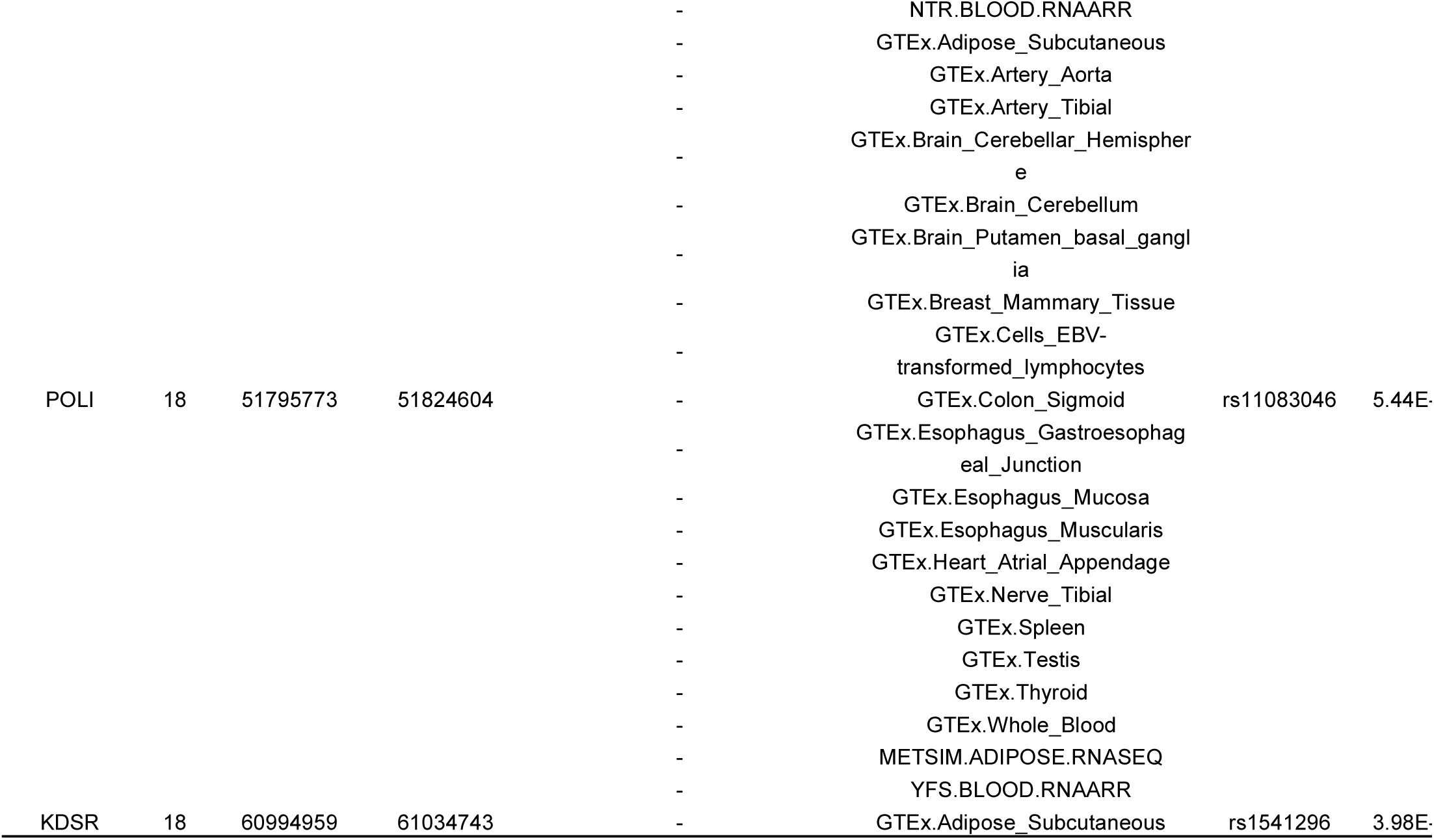
Novel risk loci. TWAS associations that did not overlap a genome-wide significant SNP (i.e. ± 1Mb transcription start site). Study denotes the original expression panel used to fit weights. P-value for TWAS computed under the null of no association between gene expression levels and PrCa risk under a Normal(0, 1) distribution. An asterisk (*) indicates associations that are nominally significant (*P* < 0.005/ 10) under a permutation test.

### TWAS replicates previously reported genes

We next sought to quantify the extent of overlapping results between TWAS and previous studies that integrated eQTL data measured in normal and tumor prostate tissues at PrCa risk regions (see Methods; see Table S7)^5^^;^ ^14^^-^^20^. We considered only autosomal, non-HLA genes which resulted in 130 previously reported genes. We found a significant overlap between reported genes, with 104/130 assayed in our study and 57/104 reaching transcriptome-wide significance in at least one of our panels (Fisher’s exact *P* < 2.2 × 10^−16^; see Tables S7-S8). For example, *MLPH* was reported in 4/8 studies. We found significant associations suggesting that decreased expression of *MLPH* in normal and tumor prostate tissue increases risk for PrCa (e.g., GTEx prostate *MLPH Z_TWAS_* = −5.80; *P_TWAS_* = 6.69 × 10^−9^; TCGA prostate *Z_TWAS_* = −6.77; *P_TWAS_* = 1.25 × 10^−11^). Predicted *MLPH* in tumor prostate remained significant under permutation, which suggests that chance co-localization with GWAS risk is unlikely (Table S2). To assess the amount of residual association signal due to genetic variation in the GWAS risk region after accounting for predicted expression of *MLPH* we performed a summary-based conditional analysis (see Methods). We found *MLPH* to explain most of the signal at its region (lead SNP *P_GWAS_* = × 10^−11^; conditioned on *MLPH* lead SNP *P_GWAS_* = 1.13 × 10^−3^; see Figure 3). Our findings are consistent with recent work that found decreased expression levels of *MLPH* to be associated with increased PrCa risk^45^. Despite previous eQTL data focusing on normal and tumor prostate tissue, we observed associations in 49 expression panels overlapping the 57 observed genes in total, underscoring earlier works demonstrating the consistency of cross-tissue cis-regulatory effects^46^.

**Figure 3.**
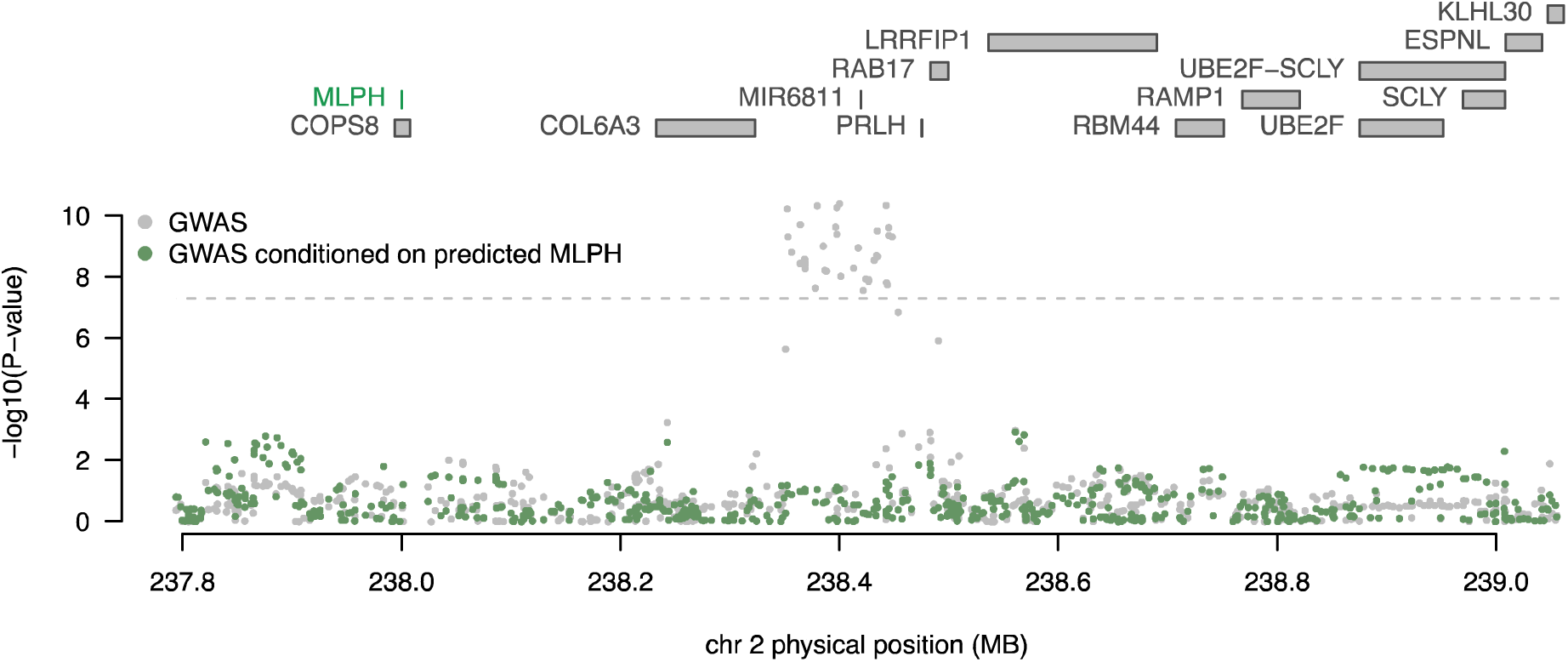
Predicted expression of *MLPH* explains majority of GWAS signal at its genomic region. Each point corresponds to the association between SNP and PrCa status. Gray points indicate the marginal association of a SNP with PrCa status (i.e. GWAS association). Green points indicate the association of the same SNPs with PrCa after conditioning on predicted expression of *MLPH* using models trained from normal prostate (GTEx) and tumor prostate (TCGA). The dashed gray line corresponds to the genome-wide significant threshold (i.e. *P* = 5 × 10^−8^). *MLPH* was discussed in previous works as a possible susceptibility gene for PrCa. Association between total expression of *MLPH* and PrCa risk was transcriptome-wide significant in normal and tumor prostate tissue.

### Bayesian prioritization pinpoints a single gene for most TWAS risk regions

TWAS genes are indicative of association and do not necessarily reflect causality (e.g., due to co-regulation at the same region). To prioritize genes at regions with multiple TWAS signals (Figure 2), we used a Bayesian formulation to estimate 90%-credible gene sets (see Methods). We found 120 unique genes across 87 non-overlapping 1Mb regions comprising our 90% credible sets (see Tables S9-S10). 71/87 credible sets contained either a single gene or the same gene in multiple tissues. The average number of unique genes per credible set was 1.38 (median 1). 27/120 prioritized genes were previously reported in eQTL analyses^5^^;^ ^14^^-^^20^, which supports the hypothesis that TWAS followed by Bayesian prioritization refines associations to relevant disease genes. For example, *MLPH* was the sole gene defining its region’s 90% credible set with a posterior probability of 94%. Similarly, *SLC22A3* (OMIM: 604842; a gene involved in polyspecific organic cation transporters^47^ and previously implicated in PrCa risk^18^) exhibited > 94% posterior probability to be causal.

### Expression and splicing events predicted in prostate tissue have largest average effect

Given the large number of significant associations observed for non-prostate tissues in our data, we wanted to quantify which tissue is most relevant for PrCa risk. We first grouped TWAS PrCa associations into prostate/non-prostate and tested for enrichment in normal and tumor prostate expression models. Predicted expression and splicing events in normal and tumor prostate made up 223/932 associations with PrCa (see Table S2) which was highly significant compared to the grouping of all other tissues (Fisher’s exact *P* = 7.8 × 10^−9^). This measure only quantifies the total amount of observed associations and neglects average association strength. Next, we computed the mean TWAS association statistic using all genes predicted from each expression reference panel (see Figure 4). We observed the largest average TWAS associations in genes predicted from normal and tumor prostate tissue, which reaffirms our intuition of expression and slice events in prostate being the most relevant for PrCa risk. We re-ranked mean associations using only genes found to be transcriptome-wide significant and observed a similar ordering with total expression in normal prostate ranked highest (average *χ*^2^ = 176.2; see Figure S11).

**Figure 4.**
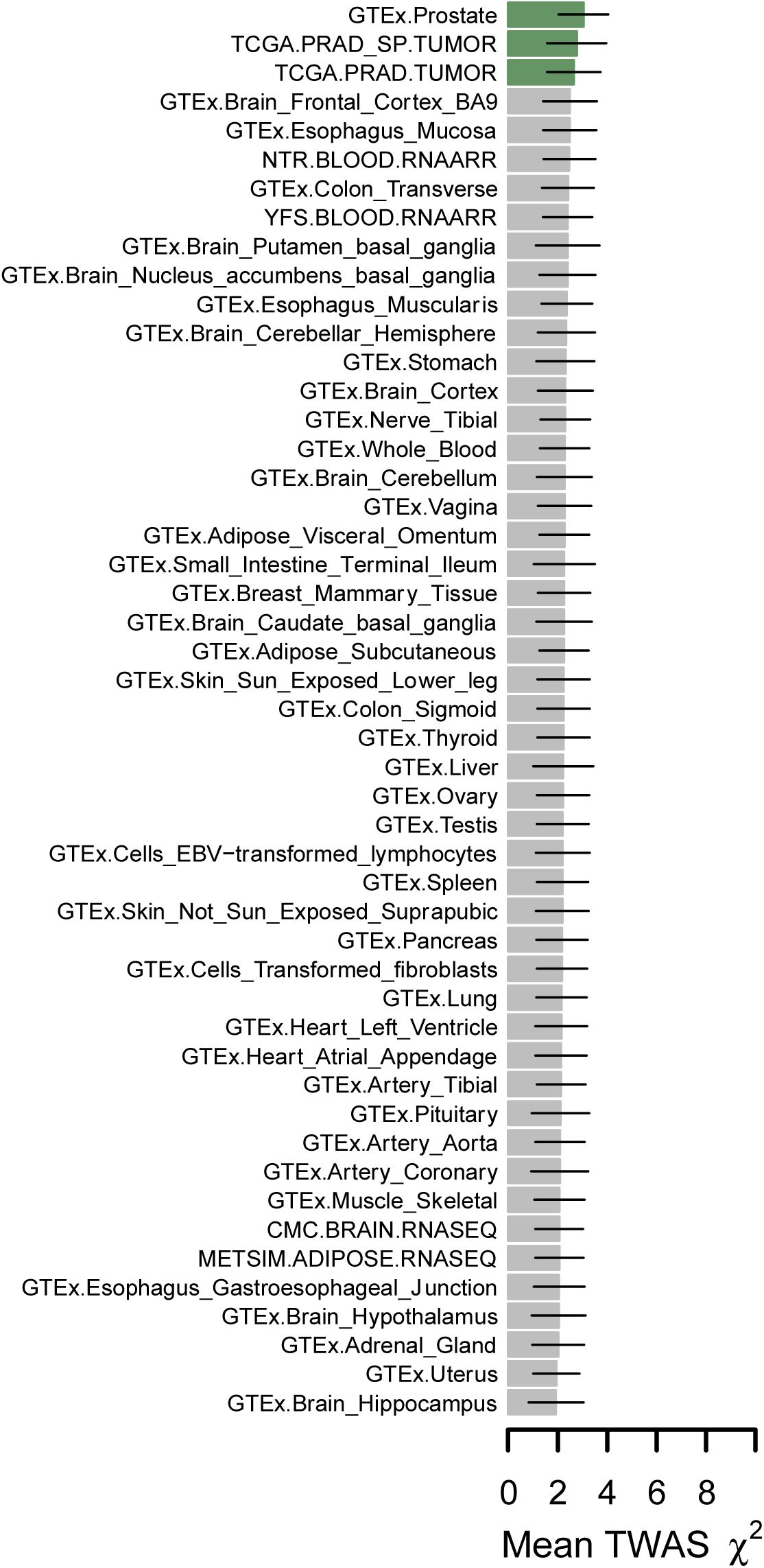
Average TWAS association statistics for genes predicted in each expression panel. Each bar plot corresponds to the average TWAS association statistic using all gene models from a given expression reference panel. Lines represent 1 standard-deviation estimated using the median absolute deviation under normality assumptions. Normal and tumor prostate tissues are marked in green.

## Discussion

Prostate cancer is a common male cancer that is expected to affect more than 180,000 men in the United States in 2017 alone^48^. While GWAS has been successful in localizing risk for PrCa due to genetic variation, the underlying susceptibility genes remain elusive. Here, we have presented results of a transcriptome-wide association study using the OncoArray PrCa GWAS summary statistics for over 142,000 case/control samples. This approach utilizes imputed expression levels and splicing events in the GWAS samples to identify and prioritize putative susceptibility genes. We identified 235 genes whose expression is associated with PrCa risk. These genes localized at 87 genomic regions, of which 9 regions do not overlap with a genomewide significant SNP in the OncoArray GWAS. We found 24 genes using predictive models for alternatively spliced introns in tumor prostate, which supports the its role in continued risk for tumor oncogenesis. A large fraction of identified genes was confirmed in earlier work, with 57 genes previously reported in eQTL/PrCa GWAS overlap studies. We used a novel Bayesian prioritization approach to refine our associations to credible sets of 120 genes with statistical evidence of causality under standard assumptions. Our results provide a functional map for PrCa risk which can be explored for follow-up and validation.

In this study, we compared our reported TWAS results with genes identified in previous works focusing on expression measured in normal and tumor prostate tissue. Several of these studies considered an eQTL and GWAS risk SNP to overlap if they are in linkage at a specified threshold. While these approaches are sound, they may be limited in statistical power for several reasons. First, if multiple local SNPs independently contribute to risk, overlap studies relying only on the top risk SNP will lose power. Second, earlier overlap studies used thresholds for association signal (i.e., GWAS *P* < 5 × 10^−8^) and linkage strength (i.e., LD > 0.5) to consider pairs of SNPs for evidence of expression influencing risk of PrCa. TWAS is largely agnostic to both issues as it jointly considers all SNPs in the region, regardless of reported GWAS association strength. However, when expression of a risk gene is regulated by a single causal SNP, we expect TWAS and earlier overlap approaches to have similar levels in power^25^.

Previous works have strongly implicated expression of certain genes in PrCa risk that were not assayed in our study (e.g., MSMB^18^^;^ ^49^) due to non-significant heritability estimates. TWAS operates by fitting predictive linear models of gene expression based on local genotype data, followed by prediction into large cohorts and subsequent association testing. Expression of genes that are not significantly heritable at current sample sizes are not included in the pipeline. This is the consequence of heritability providing an upper bound on the predictive accuracy under a linear model for genotype; therefore, if a gene has undetectable heritability at a given sample size, it will be difficult to predict using linear combinations of SNPs. To compute TWAS weights for normal prostate tissue, we used samples collected in the GTEx v6 panel (*n* = 87). Thus, our inability to detect heritable levels of gene expression can be explained due to the relatively small number of samples compared with other tissues. Indeed, previous work has shown a strong correlation between sample size in expression panels and the number of identified eGenes^27^; therefore, as sample size increases for relevant tissues, we expect the number of genes included in the TWAS framework to increase. TWAS will lose power in situations where gene expression is a non-linear function of local SNPs, or when trans (or distal) regulation is a major component in modulating expression levels.

We conclude with several caveats and possible future directions. First, while TWAS associations are consistent with models of steady-state gene expression levels altering risk for PrCa, they may be the result of confounding^25^^;^ ^26^. Imputed gene expression levels are the result of weighted linear combinations of SNPs, many of which may tag non-regulatory mechanisms driving risk and result in inflated association statistics. Second, since genes with eQTLs are common, associations may be the result of chance co-localization between eQTLs and PrCa risk. Lastly, we note recent work has extended TWAS-like methods to expose regulatory mechanisms for susceptibility genes by incorporating chromatin information^50^. An extension to our work would be to pinpoint chromatin variation regulating expression levels at identified risk genes, thus describing a richer landscape of the molecular cascade where SNP → chromatin → expression → PrCa risk.

## URLs

1000Genomes Phase3: http://www.internationalgenome.org/

Fire Hose v2016_1_28: http://gdac.broadinstitute.org/

FUSION: http://gusevlab.org/projects/fusion/

GCTA v1.26: http://cnsgenomics.com/software/gcta/

GEMMA v0.94: http://www.xzlab.org/software.html

GOseq v1.26: http://bioinf.wehi.edu.au/software/goseq/

MapSplice v2: http://www.netlab.uky.edu/p/bioinfo/MapSplice2

PLINK v1.9: https://www.cog-genomics.org/plink2/

OncoArray: https://epi.grants.cancer.gov/oncoarray/

## Funding

### CRUK and PRACTICAL consortium

This work was supported by the Canadian Institutes of Health Research, European Commission’s Seventh Framework Programme grant agreement n° 223175 (HEALTH-F2-2009- 223175), Cancer Research UK Grants C5047/A7357, C1287/A10118, C1287/A16563, C5047/A3354, C5047/A10692, C16913/A6135, and The National Institute of Health (NIH) Cancer Post-Cancer GWAS initiative grant: No. 1 U19 CA 148537-01 (the GAME-ON initiative).

We thank the following for funding support: The Institute of Cancer Research and The Everyman Campaign, The Prostate Cancer Research Foundation, Prostate Research Campaign UK (now Prostate Action), The Orchid Cancer Appeal, The National Cancer Research Network UK, The National Cancer Research Institute (NCRI) UK. We are grateful for support of NIHR funding to the NIHR Biomedical Research Centre at The Institute of Cancer Research and The Royal Marsden NHS Foundation Trust and the NIHR Biomedical Research Centre at the University of Cambridge. The Prostate Cancer Program of Cancer Council Victoria also acknowledge grant support from The National Health and Medical Research Council, Australia (126402, 209057, 251533, 396414, 450104, 504700, 504702, 504715, 623204, 940394, 614296), VicHealth, Cancer Council Victoria, The Prostate Cancer Foundation of Australia, The Whitten Foundation, Price Waterhouse Coopers, and Tattersall’s.

Genotyping of the OncoArray was funded by the US National Institutes of Health (NIH) [U19 CA 148537 for ELucidating Loci Involved in Prostate cancer SuscEptibility (ELLIPSE) project and X01HG007492 to the Center for Inherited Disease Research (CIDR) under contract number HHSN268201200008I]. Additional analytic support was provided by NIH NCI U01 CA188392 (PI: Schumacher).

Funding for the iCOGS infrastructure came from: the European Community’s Seventh Framework Programme under grant agreement n° 223175 (HEALTH-F2-2009-223175) (COGS), Cancer Research UK (C1287/A10118, C1287/A 10710, C12292/A11174, C1281/A12014, C5047/A8384, C5047/A15007, C5047/A10692, C8197/A16565), the National Institutes of Health (CA128978) and Post-Cancer GWAS initiative (1U19 CA148537, 1U19 CA148065 and 1U19 CA148112 - the GAME-ON initiative), the Department of Defence (W81XWH-10-1-0341), the Canadian Institutes of Health Research (CIHR) for the CIHR Team in Familial Risks of Breast Cancer, Komen Foundation for the Cure, the Breast Cancer Research Foundation, and the Ovarian Cancer Research Fund.

### BPC3

The BPC3 was supported by the U.S. National Institutes of Health, National Cancer Institute (cooperative agreements U01-CA98233 to D.J.H., U01-CA98710 to S.M.G., U01-CA98216 toE.R., and U01-CA98758 to B.E.H., and Intramural Research Program of NIH/National Cancer Institute, Division of Cancer Epidemiology and Genetics).

### CAPS

CAPS GWAS study was supported by the Swedish Cancer Foundation (grant no 09-0677, 11-484, 12-823), the Cancer Risk Prediction Center (CRisP; www.crispcenter.org), a Linneus Centre (Contract ID 70867902) financed by the Swedish Research Council, Swedish Research Council (grant no K2010-70X-20430-04-3, 2014-2269)

### PEGASUS

PEGASUS was supported by the Intramural Research Program, Division of Cancer Epidemiology and Genetics, National Cancer Institute, National Institutes of Health.

## Methods

### OncoArray GWAS summary statistics

Genome-wide association summary statistics for the OncoArray PrCa study were obtained from ref^29^. Summary statistics were computed using a fixed-effect meta-analysis for 142,392 total samples of European ancestry from the OncoArray (81,318/61,074 cases/controls), UK stage 1 (1,854/1,894) and UK stage 2 (3,706/3,884), CaPS 1 (474/482) and CaPS 2 (1,458/512), BPC3 (2,068/3,011), NCI PEGASUS (4,600/2,941) and iCOGS (20,219/ 20,440). The initial summary data contained association statistics for 19,726,430 variants. We filtered out summary statistics for SNPs with MAF < 0.01 and any SNPs with ambiguous alternative alleles (e.g., A→T; C→G; or vice-versa). Lastly, we kept only SNPs with rsIDs defined by dbSNP144. Our QC pipeline resulted in association statistics at 10,516,237 SNPs for downstream TWAS analyses.

### Previous studies investigating the overlap of eQTL in prostate with risk of PrCa

We collected previous studies that investigated the overlap of eQTLs in normal and tumor prostate tissue at known PrCa risk loci^5^^;^ ^14^^-^^20^. We compared TWAS statistics versus reported eQTL overlap results as aggregated in refs^14^^;^ ^15^. Across these studies, overlap of eQTLs and PrCa risk loci are computed by one of two possible methods. The first method tests known PrCa risk SNPs for association with expression levels of nearby genes/transcripts. The second method takes a two-step approach. First, genes nearby PrCa risk loci are tested for harboring eQTLs at some significance level. Next, genes with identified eQTL SNPs are tested to be in LD with known PrCa risk variants at some level (e.g., *r*^2^ > 0.5).

### Reference gene expression data sets and predictive models of expression

We downloaded the FUSION software (see URLs) along with its prepackaged weights for gene expression data. FUSION is an R package that implements the TWAS scheme described in ref^25^. Weights for gene expression measured using RNA sequencing data were obtained from the CommonMind Consortium^30^ (dorsolateral prefrontal cortex, *n* = 452), the Genotype-Tissue Expression Project^22^ (GTEx; 44 tissues; *n* = 449), the Metabolic Syndrome in Men study^32^^;^ ^33^ (adipose, *n* = 563), and The Cancer Genome Atlas (TCGA; prostate adenocarcinoma, *n* = 483)^39^. Expression microarray data were obtained from the Netherlands Twins Registry^35^ (NTR; blood, *n* = 1,247), and the Young Finns Study^31^^;^ ^34^ (YFS; blood, *n* = 1,264). All non-TCGA expression panel individuals were PrCa controls. Detailed description of quality control procedures on measured gene expression and genotype information for all non-TCGA reference panels are described in refs^25^^;^ ^27^. TCGA genotype, gene expression, and exon-junction data for 525 samples were downloaded using the Broad GDAC FireHose version 2016_1_28 (see URLs). Genotypes were imputed to the Haplotype Reference Consortium^51^ and restricted to well-imputed (INFO > 0.9) HapMap3^52^ sites. Genes (exon junctions) missing in more than half of samples were removed. RPKM and log-adjusted gene expression levels were estimated in a generalized linear model controlling for 3 gene-expression PCs and rank-normalized. We estimated alternatively spliced introns using the software MapSplice version 2 (see URLs). A total of 482 samples passed quality control procedures in both genotype and gene expression data.

We filtered genes that did not exhibit cis-genetic regulation at current samples sizes by keeping only genes with nominally significant (*P* < 0.05) estimates of cis-SNP heritability (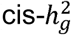), which resulted in 117,459 total tissue-gene pairs from 17,023 unique genes. We refrain from reporting genes from the HLA region due to complicated LD patterns.

To train predictive models, FUSION defines gene expression for *n* samples (***y****_GE_*) as a linear function of *p* SNPs (***X***) in a 1Mb region flaking the gene as

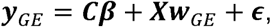

where ***w****_GE_* are the *p* SNP weights, ***Cβ*** are covariates (e.g., sex, age, genotype principal components, genotyping platform, PEER factors) and their effects, and ***ϵ*** is random environmental noise. FUSION estimated weights for expression of a gene in a tissue using multiple penalized linear models. Generally, FUSION optimizes for

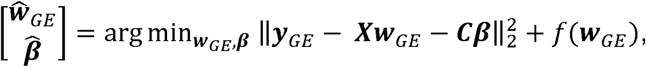

where *f*(***w****_GE_*) is a parameterized penalty function specific to each model (e.g., GBLUP^37^, LASSO, the Elastic Net). The exception to this optimization criterion is the Bayesian sparse linear mixed model (i.e. BSLMM)^38^ which fits the posterior mean for ***w****_GE_* using MCMC in the GEMMA v 0.94 software (see URLs) to obtain weights. To determine which model has the best prediction accuracy for a given gene-tissue pair, FUSION computes out-of-sample *R*^2^ by performing 5-fold cross-validation for each model. Weights from the model with the largest *R*^2^ were used to compute TWAS association statistics. We compute the normalized prediction accuracy for a gene as mim (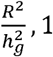).

### Cis-heritability of gene expression

FUSION reports the estimated SNP-heritability (i.e. 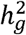) for measured gene expression levels explained by SNPs in the cis-region (1 Mb region surrounding the TSS). This is modeled under a mixed-linear model as

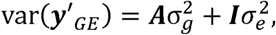

where ***y****′_GE_* is the residual gene expression after regressing out fixed-effect covariates ***C***, ***A*** is the estimated kinship matrix from SNPs in the cis-region and 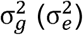 is the variance explained by the cis-SNPs (environment). SNP-heritability is then defined to be ratio of genotypic variance and total trait variance as, 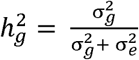. Variance parameters are estimated using the AI-REML algorithm implemented in GCTA v1.26 (see URLs) with the top 3 genotypic principal components, sex, age, genotyping platform, and PEER factors as covariates.

### Measuring cross-tissue similarity in predicted expression

We took an unbiased approach to identify susceptibility genes for PrCa by using gene expression panels measured in various tissues. To quantify how similar predicted expression levels are for the same gene across different tissues we measured the squared Pearson correlation (*R*^2^). This value represents how well predicted expression from one tissue may be used to predict expression in another tissue. To dissect similarities and differences of tissue-specific models, the ideal scenario would be to inspect effects at individual SNPs defining the models. In practice this is not possible due to predictive models not including the same set of SNPs due to QC and technological differences in the original studies. Therefore, as a proxy we predict gene expression into the 489 samples of European ancestry from 1000 Genomes^53^ and compute *R*^2^ across shared genes for pairs of tissues (see Supplementary Note).

### Transcriptome-wide association study using GWAS summary statistics

FUSION estimates the strength of association between predicted expression of a gene and PrCa (*z_TWAS_*) as function of the vector of GWAS summary Z-scores at a given cis locus ***z****_GWAS_* (i.e. vector of SNP association Wald statistics) and the LD-adjusted weights vector learned from the gene expression data ***w****_GE_* as

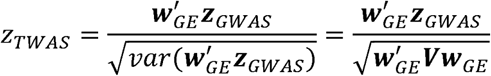

where ***V*** is a correlation matrix across SNPs at the locus (i.e. LD) and “‘” indicates transpose. A P-value for *z_TWAS_* is obtained using a two-tailed test under *N*(0, 1). In this work, we estimated ***V*** using 489 samples of European ancestry in 1000 Genomes^53^. To account for the large number of hypotheses tested, we perform a conservative Bonferroni correction at *α* = 0.05 / *M,* where *M* = 117,459 is the number of predictive models. As reported by ref^25^, there may be inflation at GWAS risk loci, due to chance co-varying of SNP effects between expression and PrCa. The same work described a permutation procedure that assesses likelihood of observing association by chance conditioned on GWAS signal. The algorithm works by permuting the eQTL weights ***w****_GE_* while keeping ***z****_GWAS_* fixed and computing *z_TWAS,perm_.* FUSION implements an adaptive procedure that stops once enough scores (i.e. |*z_TWAS,perm_*| ≥ |*z_TWAS_*|) have been observed such that the empirical null cannot be rejected at a specified level. We define novel risk regions as a flanking region around a transcriptome-wide significant gene (splicing event; *P_TWAS_ <* 4.26 × 10^−7^) that does not harbor a genome-wide significant SNP (*P_GWAS_* < 5 × 10^−8^). We consider 2Mb windows by default (i.e. TSS ± 1Mb) and show that the results are robust to the choice of window size (see Table S4).

### GWAS analyses conditional on predicted expression

To assess the extent of residual association of SNP with PrCa risk after accounting for predicted gene expression levels, FUSION estimates conditional SNP association scores using GWAS summary statistics. Namely, define ***V*** as LD for SNPs in the region, ***V****_GE_* as the correlation between predicted expression levels, and ***C*** as the correlation between SNPs and predicted expression. The least-squares estimates of ***z****_GWAS_*|***z****_TWAS_* are determined by,

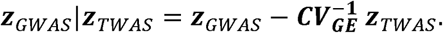

The variance of the residual association strength is given by,

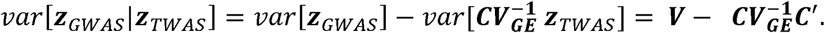

This results in the final conditional association score for the *i*th SNP as,

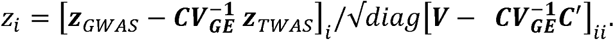

### Bayes factors and posterior inference of causal genes

Complex correlations between predicted expression levels at a given region can yield multiple associated genes in TWAS (see Figure 2). Thus, for the vast majority of risk regions it remains unclear which gene is causally influencing PrCa risk. Here we model under the assumption of a single causal gene per risk region and relying on the central limit theorem for normality, we can compute the Bayes Factor that the *i*th gene in a region is causal as,

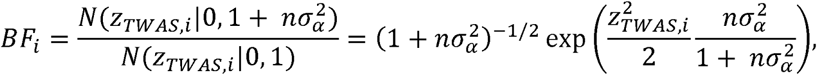

where 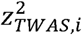 is the squared TWAS association statistic for the *i*th gene, *n* is the GWAS sample size, and 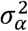 is prior effect-size variance for gene expression on PrCa risk (see Supplementary Note). This model is structurally similar in form to earlier works^54^^-^^56^ describing Bayes Factors for fine mapping SNPs at GWAS risk regions. The important distinction is that here, we formulate a Bayes Factor for genes at TWAS risk regions. The Bayes Factor for each gene quantifies the amount of evidence in favor of the causal model (*i*th gene drives risk) versus the null (*i*th gene has no causal effect). We extend individual Bayes Factors for *k* genes at a PrCa risk region to compute the posterior probability that a gene is causal as,

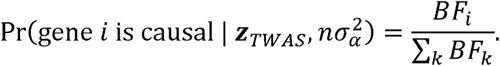

Equipped with our definition of posterior probability for each gene being causal, we define *ρ*-credible gene sets for a PrCa risk region. Formally, a set of indices *i* ∊ *I* defines a *ρ*-credible gene set if

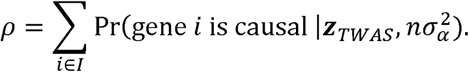

For a fixed *ρ* we optimize over k genes at a region by greedily adding genes until the total density is at least *ρ*.

To ensure that our *ρ*-credible sets are well-calibrated we performed simulations by predicting expression levels into 489 samples of European ancestry from 1000 Genomes^53^ and estimating the local correlation structure to sample TWAS Z-scores directly (see Supplementary Note). Under the assumption of a single causal gene at a risk region, we sampled TWAS Z-scores for 1000 independent regions. We then performed Bayesian prioritization at each region and computed *ρ*-credible sets for various levels of *ρ* while counting the proportion of causal genes identified across all simulations.

### Pathway analyses

To determine which pathways may be enriched with genes identified from our Bayesian prioritization approach, we used the R package GOseq^57^ which internally links gene identifiers to GO terms (GO db: 2017-09-02). We categorized all 17,023 genes into prioritized/not-prioritized and ran the analysis using custom R scripts linking GOseq. GOseq obtains P-values for overrepresented genes using the Wallenius approximation to the non-central hypergeometric distribution. We limited analysis to Gene Ontology Biological Pathways (GO:BP). GOSeq drops genes without GO categories from analysis. We observed 5,005 genes dropped from analyses resulting in 12,018 genes put forward for enrichment tests (see Table S10; Supplementary Note).

